# Pre- and post-target cortical processes predict speech-in-noise performance

**DOI:** 10.1101/817460

**Authors:** Subong Kim, Adam T. Schwalje, Andrew S. Liu, Phillip E. Gander, Bob McMurray, Timothy D. Griffiths, Inyong Choi

## Abstract

Understanding speech in noise (SiN) is a complex task that recruits multiple cortical subsystems. There is a variance in individuals’ ability to understand SiN that cannot be explained by simple hearing profiles, which suggests that central factors may underlie the variance in SiN ability. Here, we elucidated a few cortical functions involved during a SiN task and their contributions to individual variance using both within- and across-subject approaches. Through our within-subject analysis of source-localized electroencephalography, we investigated how acoustic signal-to-noise ratio (SNR) alters cortical evoked responses to a target word across the speech recognition areas, finding stronger responses in left supramarginal gyrus (SMG, BA40 the *dorsal lexicon* area) with quieter noise. Through an individual differences approach, we found that listeners show different neural sensitivity to the background noise and target speech, reflected in the amplitude ratio of earlier auditory-cortical responses to speech and noise, named as an *internal SNR*. Listeners with better *internal SNR* showed better SiN performance. Further, we found that the post-speech time SMG activity explains a further amount of variance in SiN performance that is not accounted for by *internal SNR*. This result demonstrates that at least two cortical processes contribute to SiN performance independently: pre-target time processing to attenuate neural representation of background noise and post-target time processing to extract information from speech sounds.

## 1. Introduction

Understanding speech in noise (SiN) is essential for communication in social settings. Young normal-hearing listeners are remarkably adept at this. Even in challenging SiN conditions where the speech and noise have the same intensity (i.e., 0 dB signal-to-noise ratio: SNR) and overlapped frequency components, they often recognize nearly 90% of sentences correctly (Ohlenforst et al., 2017). This suggests a surprising capacity of the auditory system to cope with noise. However, the ability to understand SiN degrades severely with increased background noise level (Ohlenforst et al., 2017), hearing loss (Harris and Swenson, 1990), and/or aging (Nabelek, 1988).

Recent studies show that normal hearing listeners show large individual differences in SiN performance (Liberman et al., 2016). The premise of this study is that by linking this variable ability for SiN perception to variation in cortical activity, we may be able to understand the neural mechanisms by which humans accomplish this ability, and this may shape our understanding of how best to remediate hearing loss.

Two broad neural mechanisms might give rise to better or worse SiN performance. First, listeners may vary in neural processing that separates the target auditory object from the mixture of sounds (i.e., similar to *external* selection processes in (Strauss and Francis, 2017). Auditory scene analysis (Bregman, 1999) processes, when occurring in parallel with auditory selective attention (Shinn-Cunningham, 2020), can inhibit the neural representation of competing sounds and enhance the neural response to attended input. This process has been conceptualized as a form of sensory gain control (Hillyard et al., 1998) or neural filtering (Obleser and Erb, 2020). Its effectiveness is often quantified as attentional modulation index (AMI), the amplitude ratio of evoked responses to background noise and target (Dai and Shinn-Cunningham, 2016; O’Sullivan et al., 2019), or the degree of neural phase-locking to the attended speech (Etard and Reichenbach, 2019; Mesgarani and Chang, 2012; Viswanathan et al., 2019). A successful sensory gain control, indicated by a positive AMI, during a SiN task will unmask the target speech from maskers, which will enhance the effective signal-to-noise ratio (SNR) in the central auditory pathway (e.g., the primary and secondary auditory cortices in the superior temporal plane: STP and the posterior superior temporal gyrus: STG).

Second, listeners might vary in neural processes for the prompt extraction of information from a speech signal (i.e., similar to *internal* selection processes in (Strauss and Francis, 2017). An inherent challenge in recognizing speech (e.g., a spoken word) is the mapping between the incoming speech cues and higher level units like words and meaning while speech unfolds rapidly over time [for review, see Weber and Scharenborg (2012), Dahan and Magnuson (2006), and Davis (2016) with references therein]. In quiet listening conditions, average young normal-hearing listeners activate a range of lexical candidates immediately at the onset of the auditory stimulus (i.e., shown by works using eye-movements in the visual world paradigm: (Allopenna et al., 1998; Dahan and Gareth Gaskell, 2007; Magnuson et al., 2007). For example, after hearing the /ba/ at the onset of *bakery*, listeners will immediately consider a range of words like *bacon*, *bathe*, or *base*, at both phonological and semantic levels. However, such rapid lexical processing develops slowly in children (Rigler et al., 2015); continuous differences in lexical processing are linked to differences in language ability (McMurray et al., 2010); and they differ in listeners with hearing loss or deteriorated acoustic-cue encoding (McMurray et al., 2019). These facts suggest that there can be individual differences in the prompt lexical processing even for *clean* speech signals.

It is as yet unclear the degree to which variation in SiN performance is related to variation in both processes (particularly in combination).

### Assessing Individual Differences in Speech Unmasking

Individual differences in the speech unmasking pathway may arise from 1) the fidelity of encoding supra-threshold acoustic features and 2) cognitive control of the domain-general attentional network. Auditory scene analysis relies on the supra-threshold acoustic features that provide binding cues for auditory grouping (Darwin, 1997). These include the spectra (Lee et al., 2013), location (Frey et al., 2014; Goldberg et al., 2014), temporal coherence (Moore, 1990; Shamma et al., 2013; Teki et al., 2011), rhythm (Calderone et al., 2014; Golumbic et al., 2013; Herrmann et al., 2016; Obleser and Kayser, 2019), and timing (Lange, 2009) of the figure and ground. The fidelity of encoding such supra-threshold acoustic features may affect the separation of target speech from background noise. Supporting this idea, previous studies have correlated the fidelity of supra-threshold acoustic cue coding to SiN understanding (Anderson and Kraus, 2010; Anderson et al., 2013; Holmes and Griffiths, 2019; Hornickel et al., 2009; Liberman et al., 2016; Parbery-Clark et al., 2009; Song et al., 2011).

Individual differences also exist in how strongly selective attention modulates cortical evoked responses to sounds (Choi et al., 2014). This suggests there may be a correlation between top-down selective attention efficacy and SiN performance. Indeed, (Strait and Kraus, 2011) reported that reaction time during a selective attention task predicts SiN performance. Similarly, studies suggest that poor cognitive control of executive attentional network predicts auditory selective attention performance (Bressler et al., 2017; Dai et al., 2018), though this has not been extended to SiN.

We can obtain a measure of the overall function of these bottom-up and top-down neural processing for speech unmasking by quantifying the amplitude ratio of early cortical auditory evoked responses to noise and target speech (similarly to the AMI concept). Here we use the N1/P2 event-related potential (ERP) components which occur with 100-300 ms latency. Previous studies showed that such ERP components are strongly modulated by selective attention but only when auditory objects are successfully segregated (Choi et al., 2013; Choi et al., 2014; Kong et al., 2015). Since those early cortical ERP components originate from multiple regions across Heschl’s gyrus (i.e., the primary auditory cortex) and its surrounding areas (e.g., posterior superior temporal gyrus) (C◻eponien et al., 1998), an efficient and collective way of indexing the neural efficiency in speech unmasking is using a scalp electroencephalographical (EEG) potential at the vertex [e.g., “Cz” of the international 10-10 system for EEG electrode montage: Koessler et al. (2009) within a limited time-window (e.g., 100-300 ms range after the stimulus onset].

### Assessing Individual Differences in Mapping Speech to Words and Meaning

To assess individual differences related to the second neural mechanism – the downstream speech information processing – we must assess a larger range of cortical regions above the auditory brainstem and cortex. Current models of speech processing suggest two distinct cortical networks (i.e., dorsal and ventral stream) that are used in parallel (Gow, 2012; Hickok and Poeppel, 2007; Myers et al., 2009; Scott and Johnsrude, 2003). The ventral stream pathway including anterior superior temporal and middle temporal gyri (STG/MTG) integrates speech-acoustic and semantic information progressively over time for the sound-to-meaning mapping (Davis and Johnsrude, 2003). The dorsal stream pathway comprising supramarginal gyrus (SMG, also known as tempo-parietal junction or TPJ) and pre-/ post-central gyri mediates the mapping between sound and articulation (Rauschecker and Scott, 2009), while inferior frontal gyrus (IFG) interacts with both pathways for lexical decision-making processes (Gow, 2012).

Studies have compared cortical responses in these areas to spoken words against acoustically-matching non-word sounds. These highlight the SMG/TPJ, MTG, and IFG in the left hemisphere as three regions that tend to exhibit more activity for words than pseudo-words (Davis and Gaskell, 2009; Taylor et al., 2013). The dual lexicon model suggested by Gow (2012) confirms the importance of those three regions by referring to left SMG and MTG as dorsal and ventral lexicons that communicate with left IFG for lexical decision making. While both SMG and MTG exhibit explicitly lexical representations, they may take complementary roles consistent with the dual stream pathway model (Gow, 2012; Hickok and Poeppel, 2007). Thus, the type of task (e.g., whether subjects are asked to make a phonological or semantic judgment) may influence the relative dominance between SMG and MTG activities during speech recognition.

Supporting the idea of broad cortical regions contributing to individual differences in speech processing, fMRI studies showed that SNR changes alter the level of neural activities across frontal, central, and temporo-parietal regions (Du et al., 2016; Vaden et al., 2015; Wong et al., 2009; Zekveld et al., 2006), while Du et al. (2016) reported the correlation between activities in fronto-central regions and speech recognition performance. However, these correlations could reflect earlier variation in speech unmasking – if auditory/attentional unmasking mechanisms are less efficient, then they could lead to differences in how strongly later regions (IFG, SMG, etc.) must work to recognize words or complete the task. Thus, it is crucial to evaluate both mechanisms simultaneously to isolate a potential role for later processes.

In addition to testing the loci of activities, it is also important to test the relative timing of activity in these pathways during speech processing. Functionally, studies using eye-movements in the Visual World Paradigm (VWP) have extensively characterized the time course of word recognition in both quiet (Allopenna et al., 1998; Dahan and Gareth Gaskell, 2007; Magnuson et al., 2007) and under challenging conditions such as noise or signal degradation (Ben-David et al., 2011; Brouwer and Bradlow, 2016; Huettig and Altmann, 2005; McMurray et al., 2017; McQueen and Huettig, 2012). Most VWP data show that, in quiet, the maximum lexical competition occurs at ~400 ms after the onset. These studies also report delayed processing under challenging conditions, but such delays do not exceed 250 ms even under the most severe degradation. This timing information can guide us when we interpret the functional implication of neural activity within certain regions; if the latency of neural activity is larger than ~400 ms, such activity may be less likely related to the online word recognition.

However, the timing of cortical activity within each pathway and the way this may be moderated by challenging listening conditions are largely unknown. This is true both within isolated regions (e.g., SMG or MTG) and across activation of broader regions (e.g., frontal lobe). This is because most of the work on speech in noise perception has been conducted with fMRI (Du et al., 2014, 2016; Wong et al., 2009; Wong et al., 2008) which has a poor temporal resolution. One study has examined evoked responses across speech processing regions using source localized EEG (Bidelman and Howell, 2016). This suggests an early response at roughly 100 ms post-stimulus in IFG. However, this study used non-sense syllables that cannot reveal lexical processes.

### The present study

The central aim of this study is to investigate the simultaneous contributions of both speech unmasking and speech recognition processes to the individual differences in SiN understanding. Our main question is whether the early-stage speech *unmasking* and later-stage *recognition* processes independently predict SiN performance, or whether the latter variable is dependent on the former. We attempted to answer this question through the combination of within- and across-subject analyses using both sensor- and source-space evoked responses in an EEG paradigm.

Subjects performed a SiN noise task in which they heard isolated consonant-vowel-consonant (CVC) English words and selected which of four orthographically presented words matched the auditory stimuli. Noise began 1 second before the speech. Two noise conditions were used: a low SNR (−3 dB) or hard condition, and high SNR (+3 dB) or easy condition. EEG was recorded from 64 electrodes while subjects performed this task, using both source- and sensor-space analyses to quantify cortical activity.

Speech unmasking and speech recognition can be distinguished by the 1) timing and 2) regional differences of neural activities evoked by both noise and target speech. Thus, we used a trial structure comprised of clearly separated events (i.e., fixed onsets of background noise and target speech) while observing time-locked neural responses to such events with electroencephalography (EEG). The degree of speech unmasking was quantified as the amplitude ratio of evoked responses to the onsets of noise and target speech measured at a vertex scalp electrode (the key scalp location for evoked responses from early auditory processes), henceforth referred to as *internal SNR*. Although the concept of *internal SNR* is similar to the attentional modulation index (Dai and Shinn-Cunningham, 2016; O’Sullivan et al., 2019), we named the index rather phenomenologically to avoid limiting the mechanism underlying the index to selective attention.

The effectiveness of later speech *recognition* processing was quantified by measuring the amplitude of evoked responses within target cortical regions. As we described, prior work has implicated a number of such regions. However, it is unclear which may be relevant for our specific task. Thus, we take a data-driven approach by first asking which post-auditory regions show *greater* activity in the *higher* SNR (easier) listening condition. This would be suggestive of a region that conducts downstream analyses once the target speech is unmasked. We looked into two regions-of-interest (ROIs): left SMG and IFG. As reviewed above, those regions activate more strongly for speech than pseudo-speech sounds; we did not consider MTG (the ventral lexicon) as our task was single CVC word identification (matched to an orthographic response). In this task, phonological discrimination (indicating the dorsal stream), rather than semantic processing, is essential.

Having quantified the contributions of each pathway, we then conducted both timing and individual differences analyses to determine their relative contribution to SiN performance.

## 2. Material and Methods

### 2.1 Participants

Twenty-six subjects between 19 and 31 years of age (mean = 22.42 years, SD = 2.97 years; median = 21.5 years; 8 (31%) male) were recruited from a population of students at the University of Iowa. All subjects were native speakers of American English, with normal hearing thresholds no worse than 20 dB HL at any frequency, tested in octaves from 250 to 8000 Hz. Written informed consent was obtained, and all work has been carried out in accordance with the Code of Ethics of the World Medical Association (Declaration of Helsinki). All study procedures were reviewed and approved by the University of Iowa Institutional Review Board.

### 2.2 Task design and procedures

We aimed to simultaneously measure SiN performance and cortical neural activity in a short (15 minute) experimental session. Sessions were kept short to avoid confounding individual differences in irrelevant psychological factors – fatigue, level or engagement ‒with individual differences in performance and processing.

Each trial (**Figure 1**) began with the presentation of a fixation cross (‘+’) on the screen. Listeners were asked to fix their gaze on this throughout the trial to minimize eye-movement artifacts. Next, they heard the cue phrase “check the word.” This enabled listeners to predict the timing of next acoustic event (the noise onset). After a 700 ms of silence, the multi-talker babble noise began and continued for 2 seconds. One second after the noise onset, the target word was heard. Finally, 100 ms after the composite auditory stimulus (noise + word) offset, four written choices appeared on the screen. The response options differed either in the initial or the final consonant (e.g., for target word *ban*, options were *than*, *van*, *ban*, and *pan;* for target word *hiss*, options included *hit*, *hip*, *hiss*, *hitch*). Subjects pressed a button on a keypad to indicate their choice and no feedback was given. The next trial began 1 second after the button press.

**Figure 1.**
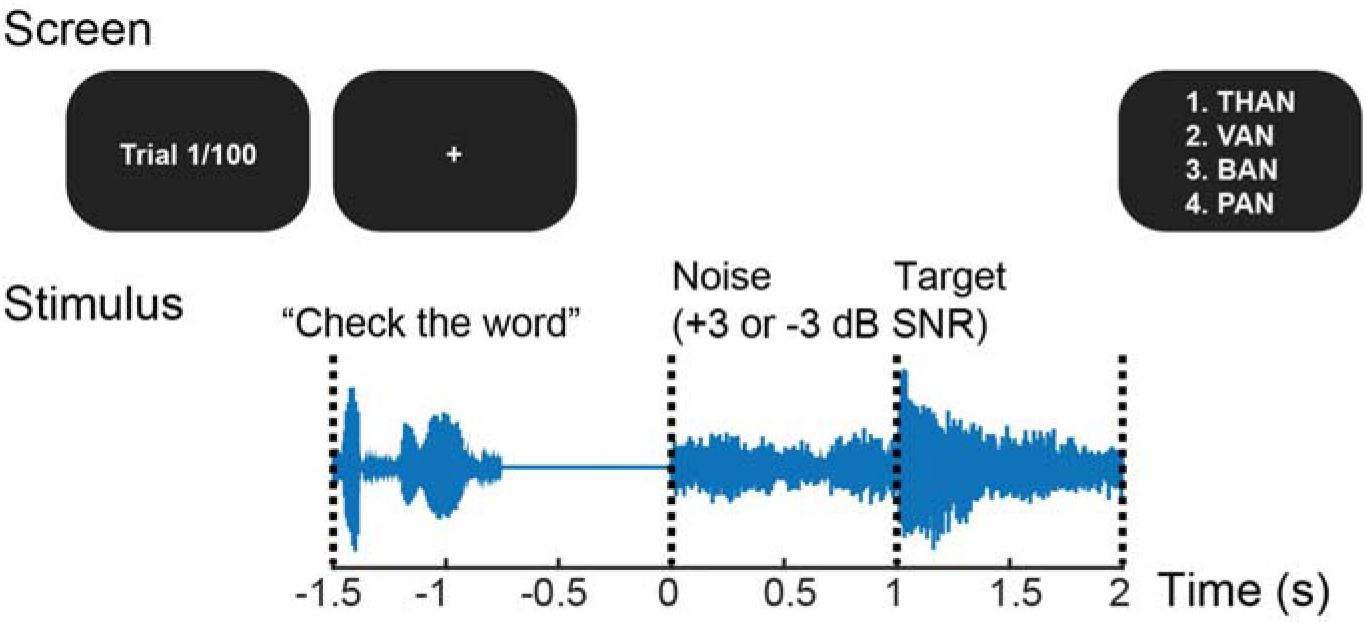
Trial and stimulus structure. Every trial starts with the cue phrase “check the word.” A target word starts 1 second after the noise onset. Four choices are given after the word ends; subjects select the correct answer with a keypad. No feedback is given. The noise level is manipulated to create high (+3 dB) and low (−3 dB) SNR conditions. Subjects complete 50 trials for each condition.

This trial structure was intended to minimize visual, pre-motor, and motor artifacts during the time of interest surrounding the auditory stimuli. The timing and intervals of auditory stimuli (i.e., cue phrase, noise, and target) were intended to derive well-distinct cortical evoked responses to the onsets of background noise and target word.

Since we were particularly interested in SMG and IFG regions that are involved in phonological and lexical processing (Gow, 2012; Hickok and Poeppel, 2007), we used naturally spoken words, rather than non-sense speech tokens used by prior EEG studies (Bidelman and Howell, 2016; Parbery-Clark et al., 2009). Target words consisted of hundred monosyllabic CVC words from the California Consonant Test (CCT) (Owens and Schubert, 1977), spoken by a male speaker with a General American accent.

Target words were always presented at 65 dB SPL. In each trial, the RMS level of noise was chosen randomly between 68 and 62 dB SPL to yield either −3 or +3dB SNR (referred to as “low SNR” and “high SNR,” respectively). Fifty words were presented at each SNR. −3 dB SNR was chosen from pilot experiments to emulate a condition yielding a mid-point performance (~65% correct) in the possible accuracy range (i.e., 25 – 100%), at which listening effort and individual differences in performance may be maximized (Ohlenforst et al., 2017). Thus, the SiN performance at −3 dB SNR condition was used for the later correlational analysis in this study. +3 dB SNR was chosen to emulate a less noisy condition from which the downstream speech recognition process will be measured for the correlational analysis.

The task was implemented using the Psychtoolbox 3 package (Brainard, 1997; Pelli, 1997) for Matlab (R2016b, The Mathworks). Participants were tested a sound-treated, electrically shielded booth with a single loudspeaker (model #LOFT40, JBL) positioned at a 0° azimuth angle at a distance of 1.2 m. A computer monitor was located 0.5m in front of the subject at eye level. The auditory stimuli were presented at the same levels for all subjects.

### 2.3 EEG acquisition and preprocessing

Scalp electrical activity (EEG) was recorded during the SiN task using the BioSemi ActiveTwo system at a 2048 Hz sampling rate. Sixty-four active electrodes were placed according to the international 10-20 configuration. Trigger signals were sent from Matlab (R2016b, The Mathworks) to the ActiView acquisition software (BioSemi). The recorded EEG data from each channel were bandpass filtered from 1 to 50 Hz using a 2048-point FIR filter. Epochs were extracted from −500 ms to 3 s relative to stimulus onset. After baseline correction using the average voltage between −200 and 0 ms, epochs were down-sampled to 256 Hz.

Since we were interested in the speech-evoked responses from frontal brain regions, we opted for a non-modifying approach to eye blink rejection: Trials that were contaminated by an eye blink artifact were rejected based on the voltage value of the Fp1 electrode (bandpass filtered between 1 and 20 Hz). Rejection thresholds for eye blink artifacts were chosen individually for each subject, and separately for the noise and target word periods. After rejecting bad trials, averages for each electrode were calculated for the two conditions to extract evoked potentials. For analysis of speech-evoked responses, we repeated baseline correction using the average signal in the 300 ms preceding the word onset.

### 2.4 Sensor-space analysis

We performed traditional sensor-space ERP analysis to investigate the effect of acoustic SNR on AC representation of noise and speech, and its individual differences. The other purpose of sensor-space analysis was to ensure the quality of data in the more familiar form before running source-space analyses.

Cortical evoked responses time-locked to the target and noise were examined and compared between high- and low-SNR conditions. EEG data were bandpass filtered from 2 to 7 Hz to capture auditory N1 and P2 components that fall into 3 – 5 Hz bands. Our first, analysis examined the N1 and P2 peak amplitude obtained from the frontal-central channels (C1, C2, FC1, FC2, FCz, Cz) and compared these measures between two SNR conditions. Both auditory N1 and P2 components were obtained at around 200 ms after the noise onset and at about 250 ms after the target word onset. After comparing the amplitude of each auditory component, we also examined the ERP envelopes by applying the Hilbert transform to the bandpass-filtered ERPs and taking the absolute value to effectively represent overall magnitude of both N1 and P2 ERP components. Then, “internal SNR” was defined as the ratio of target word-evoked ERP envelope peaks to noise-evoked ERP envelope peaks magnitude in dB scale (Equation 1). We computed this index expecting to quantify a “neural” form of an individual’s speech unmasking ability. The internal SNR is different for each subject, and is separate from the fixed external, or acoustic, SNR (here, ±3 dB).

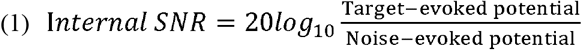

Mean activity levels were jackknifed prior to testing to assess the variance with clean ERP waveforms. In this approach, the relevant neural factors were computed for all subjects but one. This was repeated leaving out each subject in turn. The resulting statistics were adjusted for jackknifing to reflect the fact that each data point reflects N-1 subjects (Luck, 2014).

### 2.5 Source analysis

The source-space analysis was based on minimum norm estimation (Gramfort et al., 2013; Gramfort et al., 2014) as a form of multiple sparse priors (Friston et al., 2008). After co-registration of average electrode positions to the reconstructed average head model MRI, the forward solution (a linear operator that transforms source-space signals to sensor space) was computed using a single-compartment boundary-element model (Hämäläinen, 1989). The cortical current distribution was estimated assuming that the orientation of the source is perpendicular to the cortical mesh. Cross-channel EEG-noise covariance, computed for each subject, was used to calculate the inverse operators. A noise-normalization procedure was used to obtain dynamic statistical parametric maps (dSPMs) as *z*-scores (Dale et al., 2000). The inverse solution estimated the source-space time courses of event-related activity at each of 10,242 cortical voxels per hemisphere.

Following the whole-brain source estimation, we extracted representative source time courses from regions of interests (ROIs). In the present study, two predetermined ROIs were used: (1) left SMG, and (2) left pars opercularis and pars triangularis of IFG. Destrieux Atlas of cortical parcellation (Fischl et al., 2004) was used to predetermine ROIs anatomically.

Since we did not have individual structural MRI head models, it was not ideal to take the summed activity (mean or median) for all the voxels within ROIs. This is because individual difference in functional and anatomical structure of the brain may result in spatial blurring since current densities across adjacent voxels can overlap each other. Instead, representative voxels were identified for each ROI, for each SNR condition. We used a combination of previously-described methods to select voxels of interest that were used in fMRI studies (Tong et al., 2016). The voxel selection was performed by a two-step process. First, we selected voxels that exhibit greater-than-median amplitude in either (high ****or**** low SNR condition) condition. Second, cross-correlation coefficients for ERP time courses across all remaining voxels in an ROI were calculated across time, and then the mean coefficient was calculated for each voxel. The most representative voxel was defined as having the maximum mean correlation coefficient, while also being above threshold at two or more continuous timepoints based on voxel’s *p*-value, as determined using one-sample *t*-tests (Tong et al., 2016).

For the downstream statistical analyses, temporal envelopes were extracted from the within-ROI source time courses. The source time course envelopes at different ROIs were based on slightly different frequency ranges: 2-7 Hz for SMG and 1-5 Hz for IFG. This was to reflect the differences between temporal and frontal lobes in their dominant neural oscillations which create evoked activities through phase coherence (Giraud and Poeppel, 2012).

### 2.6 Statistical approaches

Once the temporal envelope of the source time course from the most representative voxel was obtained for each SNR condition, mean activity levels were compared between the two SNR conditions using paired *t*-tests. Here, we also used a jackknifing approach, and test statistics were adjusted to account for the fact that each data-point represents N-1 participants. Finally, to identify timepoints that showed a significant difference between SNR conditions while addressing multiple comparison problem, the cluster-based permutation tests were conducted (Maris and Oostenveld, 2007).

In order to identify predictors of SiN performance, sensor and source space indices of activity were used in correlation/regression analysis with SiN performance (accuracy) as the dependent variable, and the peak magnitudes of the ERP envelopes from ROIs, and internal SNR as the predictor variables. The peak magnitudes of the ERP envelopes were obtained over timepoints that showed a significant difference between high and low SNR conditions identified in the ROI-based source analysis described above. After calculating the correlation between SiN performance and these predictors, a joint contribution was tested using linear regression analysis to simultaneously examine bottom-up and compensatory related SiN performance to three factors.

## 3. Results

Our analysis started by examining the effect of SNR on task performance (accuracy and reaction time). This was intended to document that noise manipulation had the expected effect. Next, we examined the effect of SNR on both the magnitude and timing of neural activity. This was done first in the sensor-space, using the auditory N1/P2 components and other measures to examine primary auditory pathways. Next, this was done in the source-space to examine the compensatory role of IFG, to evaluate whether IFG effects were early enough to play a role in speech perception, and to test hypotheses about SMG. Finally, we turn to our primary analysis: a regression testing the unique contributions of each pathway to individual differences in SiN performance. Original raw and processed data of the present study are available at Mendeley Data (http://dx.doi.org/10.17632/jyvythkz5y.1).

### 3.1 SiN performance

There was a large variance in performance among participants. This was observed in both the high SNR condition (accuracy: mean = 80.64%, SD = 7.81%, median = 83.01%; reaction time: mean = 1.53 s, SD = 0.32 s, median = 1.55 s) and the low SNR condition (accuracy: mean = 68.21%, SD = 8.92%, median = 70.37%; reaction time: mean = 1.70 s, SD = 0.36 s, median = 1.69 s). There was a significant effect of SNR on both accuracy (*t*(25) = 6.99*, p* < 0.001, paired t-test) and reaction time (*t*(25) = −3.81*, p* < 0.001, paired t-test) (**Figure 2A**). Reaction time and accuracy were correlated in the high SNR condition (**Figure 2B**, *r* = −0.50, *p* = 0.009), but not in the low SNR condition (**Figure 2C**, *r* = −0.19, *p* = 0.34).

**Figure 2.**
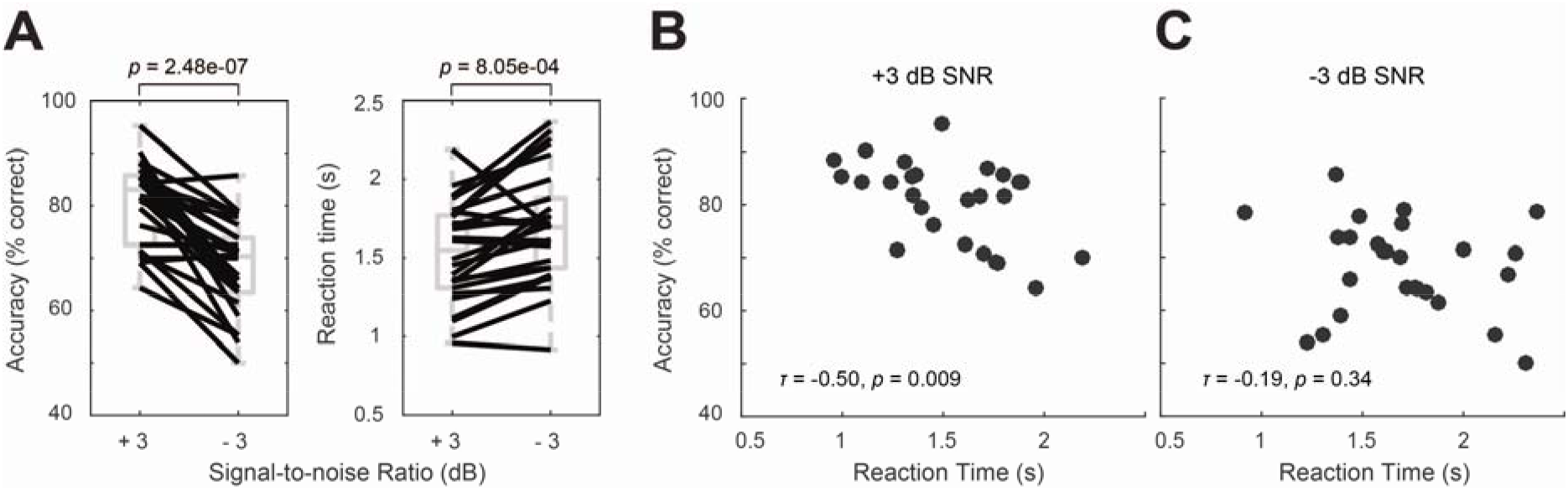
Behavioral results. **A.** Summary of behavioral performance for the two conditions (+3 and −3 dB SNR). Boxes denote the 25^th^ – 75^th^ percentile range; the horizontal bars in the center denote the median; the ranges are indicated by vertical dashed lines. Solid lines connect points for the same subject in different conditions. **B.** Average accuracy as a function of reaction time in +3 dB SNR condition. **C.** Average accuracy and reaction time in −3 dB SNR condition.

As a whole, these results validate that the SNR manipulation was sufficient to create differences in speech perception. The negative correlation between the accuracy and reaction time may demonstrate the redundancy between individual differences of accuracy and difficulty.

### 3.2 Auditory N1-P2 components in the sensor space

We next examined the N1 and P2 peak amplitude for both noise- and target word-evoked responses. This was done to estimate the contribution of primary auditory pathways and the effect of noise. These measured were compared between high- and low-SNR conditions.

We started with the noise-evoked responses (i.e., peak ERP components arising ~0.2 seconds following the noise onset at 0 s). Here, both N1 (*t(25)* = −2.64, *p* = 0.014) and P2 amplitudes (*t(25)* = −3.82, *p* < 0.001) were greater in the low SNR condition. In contrast, target-word evoked responses (i.e., peak ERP components arising ~0.25 seconds following the target-word onset at 1 s) showed the opposite pattern with stronger N1 (*t(25)* = 1.87, *p* = 0.073) and P2 amplitude (*t(25)* = 2.41, *p* = 0.023) in the high-SNR condition (**Figure 3**).

**Figure 3.**
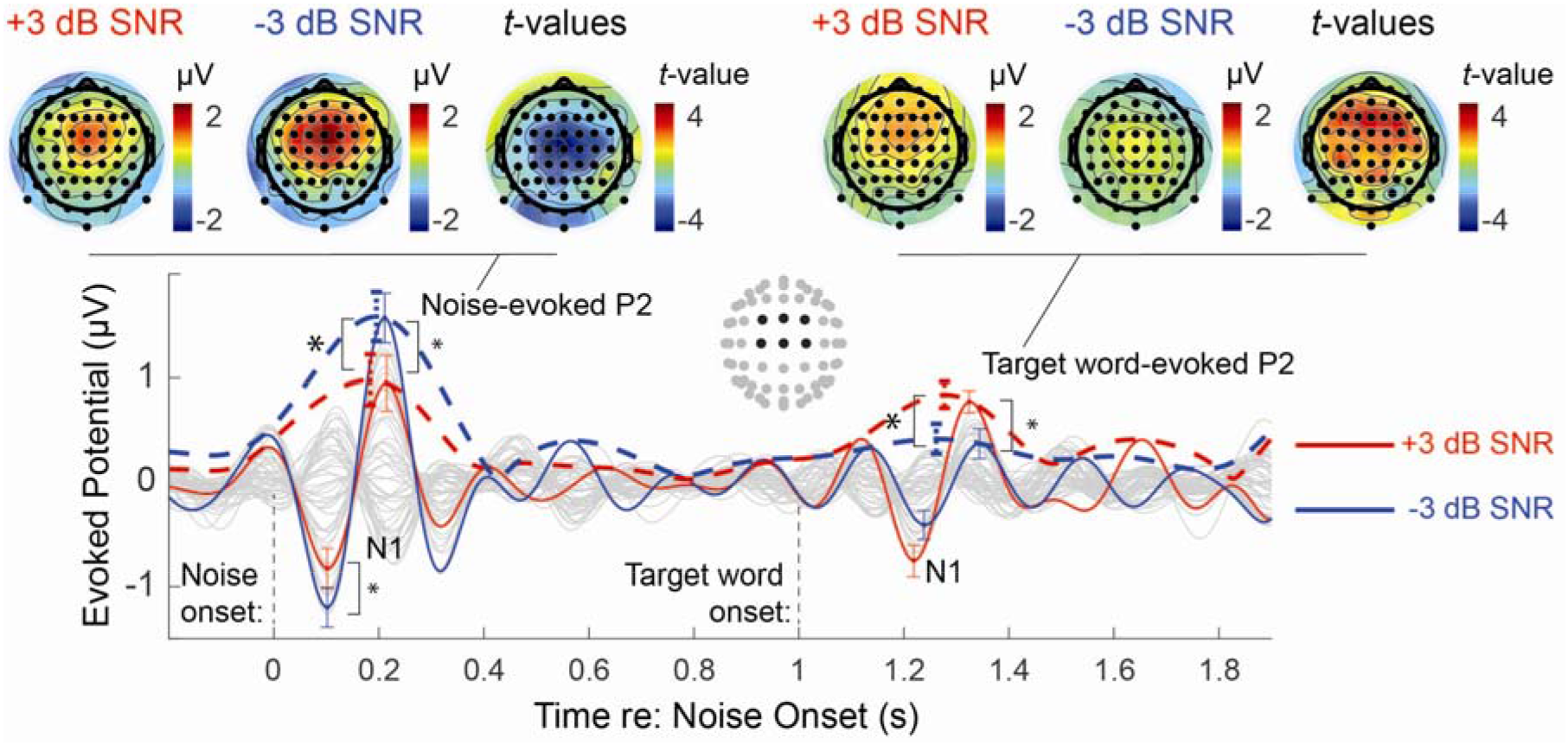
Sensor-level event-related potential (ERP) with the topographical layout and cortical maps. The time course of the auditory ERP and its envelope, with the standard error of the mean (±1 SEM) at the peak amplitude (red color: +3 dB SNR, blue color: −3 dB SNR). An asterisk shows a significant difference in the amplitude between +3 and −3 dB SNR conditions (paired *t*-test). Top panels show peak P2 amplitudes of all electrodes in topographical layouts. The *t*-values from paired *t*-tests between two SNR conditions are also shown as topographies.

We also examined the ERP envelopes in the time region of both the auditory N1 and P2. Again in the noise-evoked time region, we saw a greater response in the low SNR condition (*t(25)* = −3.95, *p* < 0.001). However, also as in the N1/P2 analysis we saw larger word-evoked response in the high SNR condition (*t(25)* = 2.37, *p* = 0.026) (**Figure 3**). Then we calculated the internal SNR, the magnitude ratio of the noise and target-word related ERP envelopes, and saw a greater internal SNR in the high SNR condition (*t(25)* = 2.53, *p* = 0.018).

The topographical layout of these effects is shown in **Figure 3** (top panels) at the time of the peak P2 (which showed a significant overall effect of noise level) for both noise- and word-evoked response. T-values from paired t-tests on peak P2 amplitudes at all electrodes between SNR conditions were also represented in topographies. These show a broad-based effect that is roughly centered at frontal-central channels for both the noise onset and speech onset (though patterns for the speech evoked P2 were more distributed along the scalp). This justifies our use of frontal-central channels for sensor-space analyses. The significant difference in auditory ERP envelopes and in the N1/P2 components according to the noise level supports our use of the internal SNR as an index of individual ability to modulate representations of target speech relative to noise in the regression analysis.

### 3.3 The effect of SNR on cortical activity

Next, we conducted parallel analyses in source space to assess cortical activity through speech processing regions. We converted sensor-space EEG signals to whole-brain source time courses to localize the effects of SNR on evoked responses within targeted ROIs. Within left SMG, the cluster-based permutation test (Maris and Oostenveld, 2007) revealed that the high SNR condition evokes significantly greater activity than the low SNR condition from 270 to 340 ms (*p* = 0.0020) (**Figure 4A left**). High-SNR peak amplitude is found at 309 ms. Such a significant SNR effect was not found in the left IFG. The maximum magnitude of the grand average low-SNR evoked response (dSPM) is at 770 ms (**Figure 4B left**). Source time courses in the right SMG and IFG are shown in **Figure 4A and B** (the right panels) for visual comparisons.

**Figure 4.**
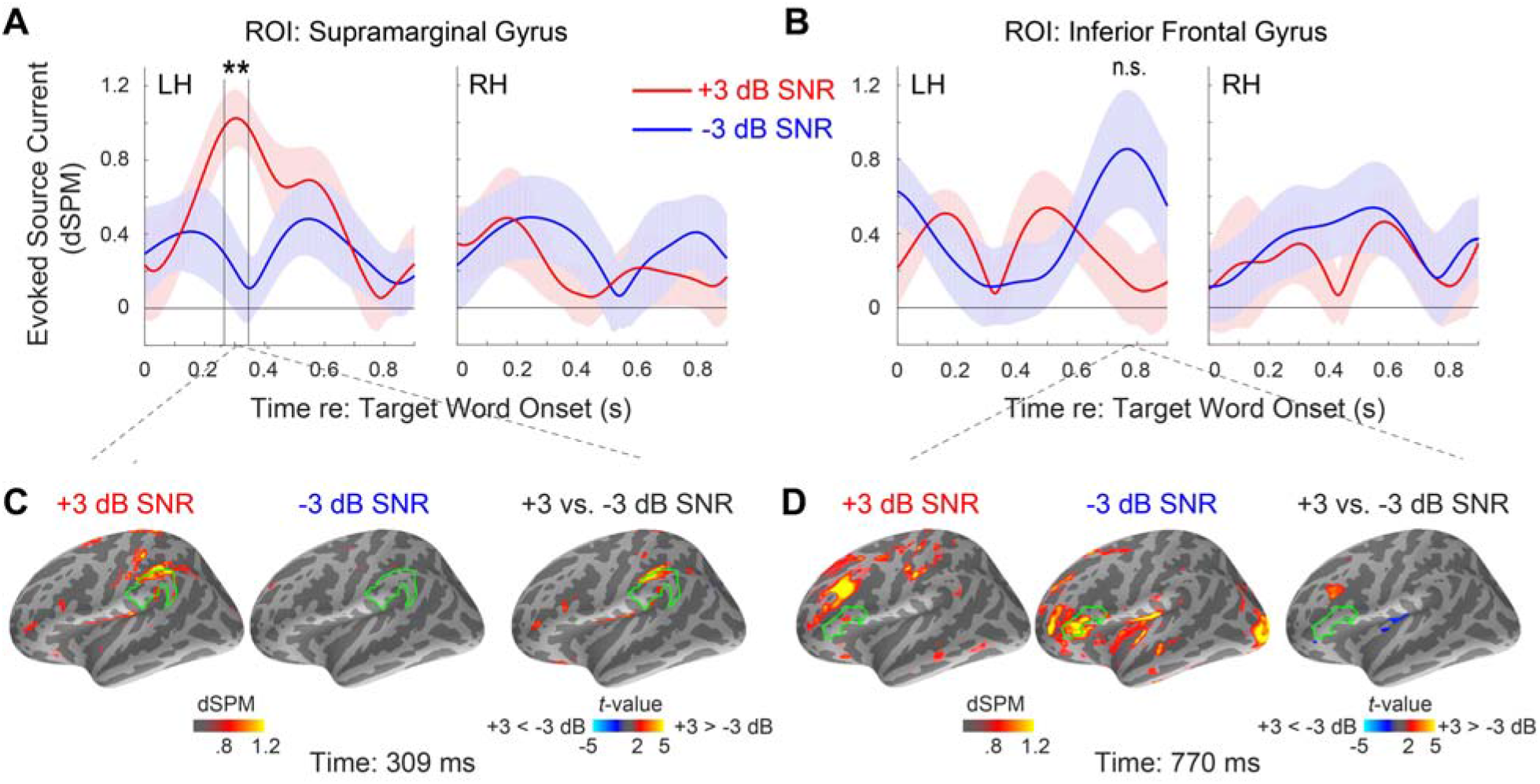
Region-of-interest (ROI) based source analysis. **A and B.** The time course of the event-related potential (ERP) envelope, with the standard error of the mean (±1 SEM), obtained at representative voxels for two ROIs in the left hemisphere (“LH”): supramarginal gyrus (SMG), and the pars opercularis and triangularis of the inferior frontal gyrus (IFG), respectively, in each SNR condition (red color: +3 dB SNR, blue color: −3 dB SNR). The ERP envelope time courses in the corresponding regions in the right hemisphere (“RH”) are also shown for visual comparisons. Asterisks show the timing of a significant difference between +3 and −3 dB SNR conditions (cluster-based permutation test, *p* = 0.0020) at the left SMG. **C and D.** Whole-brain maps showing statistical contrasts (*t*-values obtained from post-hoc paired *t*-tests between the two SNR conditions) of source activation at each voxel at the peak timepoint of the grand-average source time course of each ROI.

Single time-point evoked-current estimates are shown for the peak SMG activity time (i.e., 309ms, **Figure 4C**) and IFG peak time (770ms, **Figure 4D**) on the whole left-hemisphere cortical surface. At 309ms, post-hoc paired *t*-tests on all the left-hemisphere voxels reveal an area near left SMG that shows greater evoked responses in the high SNR condition. This supports a significant role for SMG in SiN processing in this task. In contrast, at 770ms, no voxel was found within IFG that shows significant differences between high-vs. low-SNR conditions. This confirms our timecourse analyses of IFG, suggesting it does not play a large role and that the trend that was observed is not broadly seen across voxels.

To demonstrate the timing of SMG activity compared to the timing of phonological events, the webMAUS (Kisler et al., 2017) was used to identify the boundaries between the first and second, and the second and third phonemes in each of the 100 stimuli. A histogram of these acoustic time points is shown in **Figure 5**. This confirms that the peak of evoked activation in SMG (i.e., ~309 ms, denoted by a dashed vertical line) occurs within the time course of target words before the final phoneme is presented.

**Figure 5.**
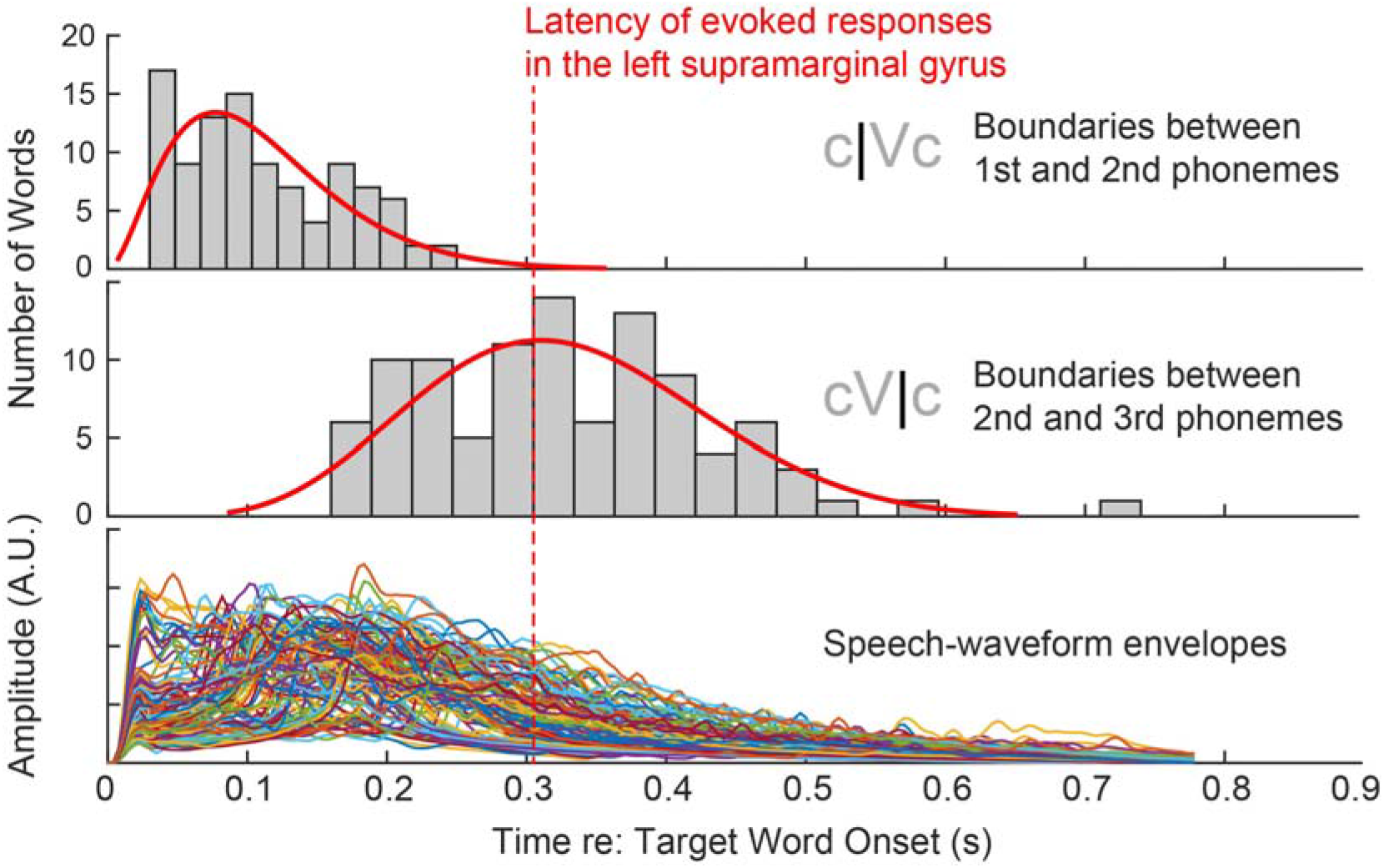
Top and second panel show histograms of boundaries between phonemes of each stimulus. The third panel shows superimposed temporal envelopes extracted from waveforms of the 100 words.

### 3.4 Individual differences in internal SNR predict SiN performance

To address our primary research question, which was to evaluate the simultaneous contribution of speech unmasking and recognition processes to SiN performance, we conducted a linear regression analysis in which internal SNR and SMG activation were used as independent variables. SiN performance in low SNR condition, which showed larger variance (high SNR SD = 7.81%; low SNR: SD = 8.92%), was used as the dependent variable. We extracted the internal SNR from the low SNR and the SMG activity from the high SNR condition, as we expected that the internal SNR captures how well listeners unmask speech from the noisy background while the SMG activity reflects the processing of relatively clean speech signal. As expected, those two metrics extracted from different trials did not show a correlation (*r* = −0.04, *p* = 0.863, the left panel of **Figure 6D**). Both internal SNR (*t*(23) = 3.35, *p* = 0.003) and SMG activity (*t*(23) = 2.29, *p* = 0.031) were significant predictors of SiN performance (**Figure 6A**). The linear combination of those predictors accounted for a large proportion of the variance (*r* = 0.64, *p* = 0.00043, **Figure 6B**).

**Figure 6.**
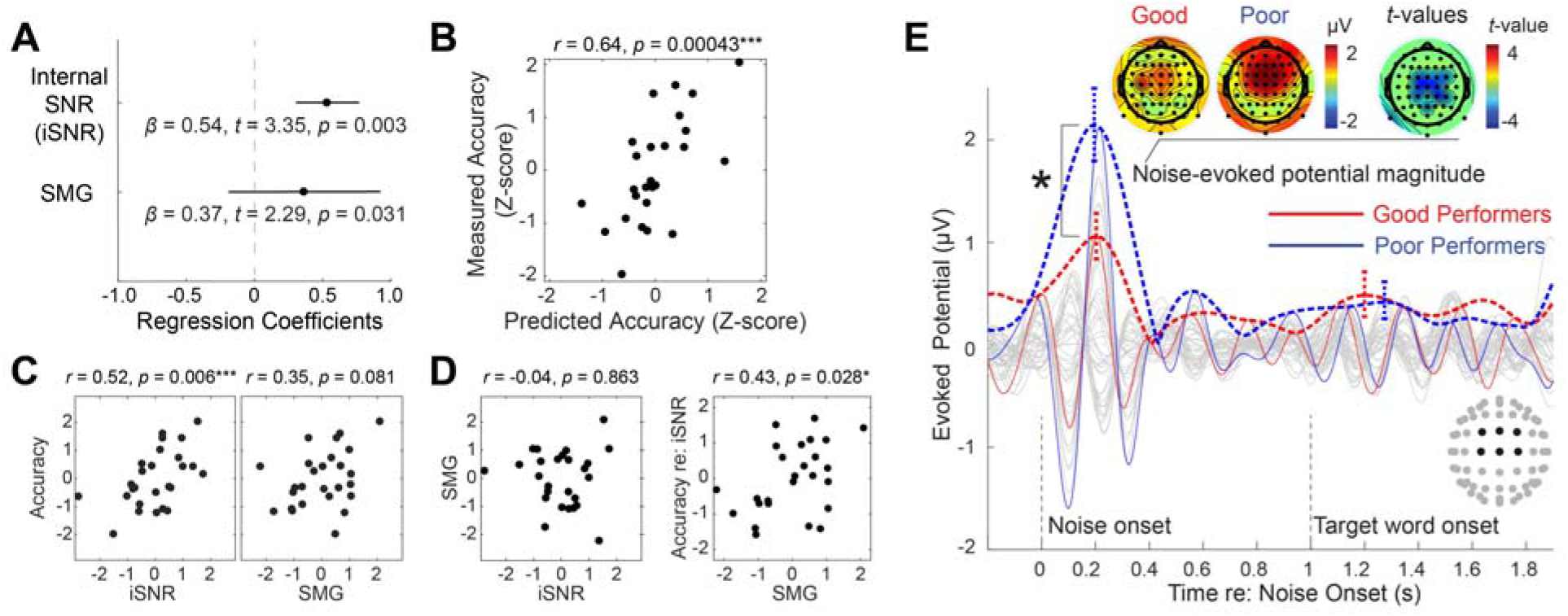
Individual differences in speech-in-noise processing. **A**. Regression coefficients and their standard errors. **B**. A scatter plot showing the relationship between predicted and measured accuracy in −3 dB SNR condition. **C**. Post-hoc correlation analyses: Raw correlations between each independent variable and the dependent variable. **D**. Left: Relationship between independent variables shows no correlation between internal SNR and evoked source current at the left supramarginal gyrus (SMG). Right: Semi-partial correlation between SMG evoked source current and the residual of accuracy after regressing out internal SNR. **E**. The time course of the auditory event-related potential and its envelope, with the standard error of the mean (±1 SEM) at the peak magnitude in −3 dB SNR condition (red color: good performers, blue color: poor performers). An asterisk shows a significant difference in the magnitude between two groups (two-sample *t*-test).

**Figure 6C and D** show results from post-hoc correlational analyses. Internal SNR showed a significant correlation with accuracy, while SMG activation did not (despite its significant contribution to the model). There was no correlation between internal SNR and SMG activation, as described above. A semi-partial correlation between SMG activation and the residual of accuracy after regressing out internal SNR was significant, which confirmed that the SMG activation accounted for an extra amount of variance in SiN performance (the right panel of **Figure 6D**). This suggests that in order to identify the contribution of downstream recognition areas like SMG, models must account for the contribution of earlier upstream speech-unmasking processes.

To visualize the contribution of internal SNR to SiN performance, **Figure 6E** showed evoked response differences between good and poor performers (based on a median split on the low-SNR condition accuracy). This reveals dramatic differences in the magnitude of noise onset-related potentials: despite the same physical noise level for each group, good performers exhibited less strong evoked response to noise onset, measured by the envelope peak magnitude within N1-P2 time range in the frontal-central channels (two-sample t-test *t*(24) = −2.60*, p* = 0.016). In contrast, the word-evoked ERP envelope did not show a significant difference between the two groups (*t*(24) = 0.21*, p* = 0.84, two-sample *t*-test). This suggests that the neural mechanism underlying the internal SNR variance is the suppression of noise (rather than the enhancement of the target).

## 4. Discussion

We investigated neural correlates of SiN performance in young normal hearing adults. Previous correlational studies focused on the contributions of either acoustic encoding fidelity (Anderson and Kraus, 2010; Anderson et al., 2013; Holmes and Griffiths, 2019; Hornickel et al., 2009; Liberman et al., 2016; Parbery-Clark et al., 2009; Song et al., 2011) or the degree of speech/language network recruitment (Du et al., 2016) to the SiN performance. However, the relative importance of each process has remained unclear. We showed that 1) how well the listener suppresses background noise *before* hearing the target speech and 2) how strongly the listener recruits temporo-parietal network *while* the speech signal is received contribute to the SiN performance independently. Combining those two factors explained about 40% of the variance in SiN performance.

Our results have both theoretical and clinical implications. Theoretically, our individual difference approach revealed at least two neural subsystems involve during SiN processing: sensory gain control and post-auditory speech recognition processing. Clinically, our results suggest that a relatively short (~15 minutes) SiN-EEG paradigm can assess crucial neural processes for SiN understanding.

### Internal SNR: A measure of pre-speech processing for speech unmasking

The first among the two crucial processes – how well the listener suppresses background noise – was indexed as “internal SNR,” the ratio of noise-to target word-evoked cortical responses. This process can be understood as pre-target cortical activity, appearing as an enhanced neural representation of the target sound (the speech) and suppressed neural representation of ignored stimuli (the noise).

### What is the source of variation in the internal SNR?

Such responses could reflect auditory selective attention, which shows a similar pattern in previous studies (Hillyard et al., 1973; Hillyard et al., 1998; Mesgarani and Chang, 2012). In the present study, good performers showed significantly weaker noise-evoked responses at frontal-central channels (around Cz), compared with poor performers, approximately 200 ms after the noise onset (**Figure 6E**). Decreased auditory responses to background noise in good performers are compatible with the presence of a sensory gain control mechanism (Hillyard et al., 1998) which may happen in multiple sub-regions in STP and posterior STG. The variation in the sensory gain control may originate from multiple factors. It may reflect the acuity of encoding spectro-temporal acoustic cues from speech and noise, or grouping of such acoustic cues for auditory object formation (Moore, 1990; Shamma et al., 2013; Teki et al., 2011). How robustly the low-frequency neural oscillations (e.g., theta and delta) are phase-locked to the acoustic temporal structure of the stimuli (Etard and Reichenbach, 2019) may also contribute to the variation, as the neural phase-locking relies on the encoding of acoustic cues (Ding et al., 2014) and the prediction of temporal structure in speech rhythm (Ding et al., 2016). Since our experiment provided fixed timing of noise and target word onsets, the neural phase-locking based on predicted timing (Arnal and Giraud, 2012) could occur and contribute to the internal SNR. The variation may also reflect endogenous mechanisms for active suppression of background sounds along with neural enhancement of foreground sounds (Shinn-Cunningham and Best, 2008). It was not our goal to disentangle the sources of variation in sensory gain control. Rather, we aimed to quantify the effectiveness of sensory gain control by our unique trial structure that enables clear distinction of evoked responses to noise and target speech, and to test how the internal SNR predicts later speech processes and behavioral accuracy. In this regard, we found a significant correlation between accuracy and the relative magnitude of word- and noise-evoked potentials.

### Evoked amplitude in SMG: the neural marker of effective and prompt lexical processing

While the computation of internal SNR was pre-specified, we had an open plan for extracting a representative neural factor to capture post-auditory speech recognition. To explore such neural markers, we added a 6-dB higher SNR condition and asked which region or regions showed increased activity within a reasonable (200 – 500ms) time range. We investigated two ROIs: left SMG and left IFG. As left SMG showed increased evoked response to target speech in the less noisy condition at ~300 ms after the target onset, the peak evoked amplitude in left SMG measured in the high SNR condition was used as the second independent variable in the regression analysis.

### Functional interpretation of SMG activity

Previous studies have suggested that spoken-word recognition occurs via a process of dynamic lexical competition as speech unfolds over time. The VWP studies reported that, for many words, this competition maximizes around 3-400 ms after word onset (Farris-Trimble and McMurray, 2013; Huettig and Altmann, 2005). In significantly challenging conditions (high noise), however, lexical processing can be delayed about 250 ms until most of the word has been heard (Farris-Trimble et al., 2014; McMurray et al., 2017), which may minimize competition. The latency of SMG activity that lied between the second and the third phonemes (see **Figure 5**) in the high SNR condition aligns well with the timing of lexical competition found from the VWP studies, which may suggest that the SMG activity makes a neural substrate of immediate lexical access (Farris-Trimble et al., 2014; McMurray et al., 2017), consistent with Gow (2012). This immediacy was observed when speech sounds were relatively clean (high SNR), and it does not appear in previous EEG studies using non-word synthesized phonemes (Bidelman and Dexter, 2015; Bidelman and Howell, 2016).

After the contribution of speech unmasking (i.e., internal SNR) is regressed out, the SMG evoked amplitude in the cleaner condition predicted the residual of SiN performance (the right panel of **Figure 6D**). This indicates that changes in SMG activity may be an independent factor predicting speech recognition performance, rather than the outcome of pre-speech sensory gain control processing.

### Limitation of the current study

In the present paper, we exhibited evoked responses only. Although our results demonstrated how this simple and traditional EEG analysis successfully predicted SiN performance, future studies may pursue further understanding of SiN mechanisms by adopting extended analyses such as induced oscillation (e.g., Choi et al. (2020)) and connectivity analyses.

Our correlational result is limited to young normal hearing listeners where variance does not come from hearing deterioration and aging. This study does not predict how much of the variance can be explained by the combination of internal SNR and SMG activity in the population with larger age ranges. For example, Tune et al. (2020) reported that no correlation is found between neural factors and behavioral success in a large cohort of aged listeners.

We chose −3 dB as the main SNR from which the dependent variable for the regression analysis was extracted. Although we claim that the −3 dB SNR provides the most representative condition where the individual difference in performance is maximized, we do not claim that our main findings can be generalized to other SNR conditions.

### Methodological advances and justifications for source time course analysis

Our approach to identifying a single voxel within an ROI deserves a particular discussion. Identification of the representative voxel of an ROI is a problem common to EEG source analysis, fMRI, and other functional brain imaging studies. Many relevant neuroimaging analysis approaches have been described, including univariate, multivariate, and machine learning; however, most of these are intended for the identification of regions of interest or functional connections from a whole-brain map. Drawbacks of this type of whole-brain analysis include the need for strict multiple comparisons correction and, therefore, decreased statistical power.

Using strong a priori hypotheses to generate regions of interest allowed us to circumvent these issues, but still requires identification of representative voxels within our regions of interest. Favored approaches generally require identification of peak activity within an ROI (Tong et al., 2016). However, to avoid the assumption that choosing peak activity implies, we opted instead to choose the voxel that has the maximum average correlation to every other voxel within the ROI. In the present study, we chose not to constrain the location of the voxel of interest within an ROI for each condition. Because our anatomic resolution is unlikely to be at the voxel level, we elected to choose a different representative voxel for each condition, unconstrained by the location of the representative voxel from other conditions.

## 5. Conclusion

We found that better speech unmasking in good performers modulated the ratio of cortical evoked responses to the background noise and target sound, which effectively changed SNR internally, resulting in better performance. We also found that clean, intelligible speech elicits early processing at SMG, which explained an extra amount of variance in SiN performance. These findings may collectively form a neural substrate of individual differences in SiN understanding ability; the variance in SiN perception may be a matter of both primary processes that extract the signal from noise and later speech recognition processes to extract lexical information from speech signals promptly.

## Acknowledgments

This work was supported by Department of Defense Hearing Restoration and Rehabilitation Program grant awarded to Choi (W81XWH1910637), NIH T32 (5T32DC000040-24) awarded to Schwalje, and NIDCD P50 (DC000242 31) awarded to Griffiths and McMurray. The authors declare no competing financial interests.

